# Woody plants constructing tundra soils

**DOI:** 10.1101/789743

**Authors:** Julia Kemppinen, Pekka Niittynen, Anna-Maria Virkkala, Konsta Happonen, Henri Riihimäki, Juha Aalto, Miska Luoto

**Author notes:** **Author contributions** JK conceived the research and together with PN, A-MV, HR, and ML designed the study setting. JK, PN, and A-MV performed the field research. PN provided the soil temperature calculations, A-MV performed the soil organic carbon stock calculations, and HR provided the calculations on LiDAR-based variables. JK, with support from KH, analysed the data. JK, with support from all of the co-authors, wrote the paper.

## Abstract

In tundra, woody plants are expanding towards higher latitudes and altitudes due to increasingly favourable climatic conditions. Their expansion may also occur through increases in the coverage and height of the plants. These shifts may cascade further across the ecosystem, such as in the foundations of tundra: that is, in the soils. Yet, little is known about the effects woody plants have on local soil conditions. Here, we examined if the coverage and height of woody plants affect the growing-season soil moisture and temperature as well as soil organic carbon stocks. We carried out a field observation study in a dwarf shrub–dominated tundra and built a hierarchical model. We found that, after controlling for other possible factors influencing woody plants and soil conditions (namely, topography, snow, and the overall plant coverage), the coverage of woody plants inversely correlated with all three soil conditions. Yet, we found no link between the woody plant height to the soil variables. This indicates that woody plants affect local soil conditions in various ways, depending upon whether their expansion occurs though the growth of coverage or their height. Nevertheless, woody plants likely alter the very ground of the entire tundra system and feedback into the global climate system through the water, energy, and carbon cycles of tundra.

## Introduction

In tundra regions, climate change has led to an increase in woody plant dominance through, for instance, shrub expansion and shrubification (Myers-Smith et al., 2011). This results from increasingly favourable environmental conditions, such as rising growing-season temperatures, thicker snow cover, and increasing winter precipitation (Carrer, Pellizzari, Prendin, Pividori, & Brunetti, 2019; Hallinger, Manthey, & Wilmking, 2010). The expanding distribution and increasing coverage and height of woody plants carry the potential of affecting tundra ecosystems through species composition and disturbance frequency including fire activity (Jeffers, Bonsall, Watson, & Willis, 2012; Mod & Luoto, 2016; Scharnagl, Johnson, & Ebert-May, 2019). Moreover, taller woody plants may decrease the tundra albedo when their top branches reach the snow surface during winter. In spring, this may further result in earlier snowmelt substantially amplifying surface warming (Chapin et al., 2005; Rydsaa, Stordal, Bryn, & Tallaksen, 2017).

Plants are the key components at the interface between atmospheric and soil processes (Loranty et al., 2018; Parker, Subke, & Wookey, 2015; Robinson et al., 2019). Changes in tundra ecosystems are closely linked to the global climate system through water, energy, and carbon cycles (Cahoon et al., 2012). Through the processes in these cycles, the expansion of woody plants, the largest plant life form of tundra, is likely to lead to cascading effects across the tundra biome and beyond (Chapin et al., 2005; Myers-Smith et al., 2011). Yet, a gap in our knowledge exists regarding how woody plants modify soil conditions on the very substrate upon which they grow. Vegetation can, in general, influence the amount of vital resources and other soil conditions, such as the soil moisture, temperature, and carbon stocks, of tundra ecosystems via multiple pathways (Figure 1) (Aalto, le Roux, & Luoto, 2013). Thus, vegetation can be viewed as a key component in soil formation, although it remains insufficiently included in tundra soil research (Robinson et al., 2019).

**Figure 1.**
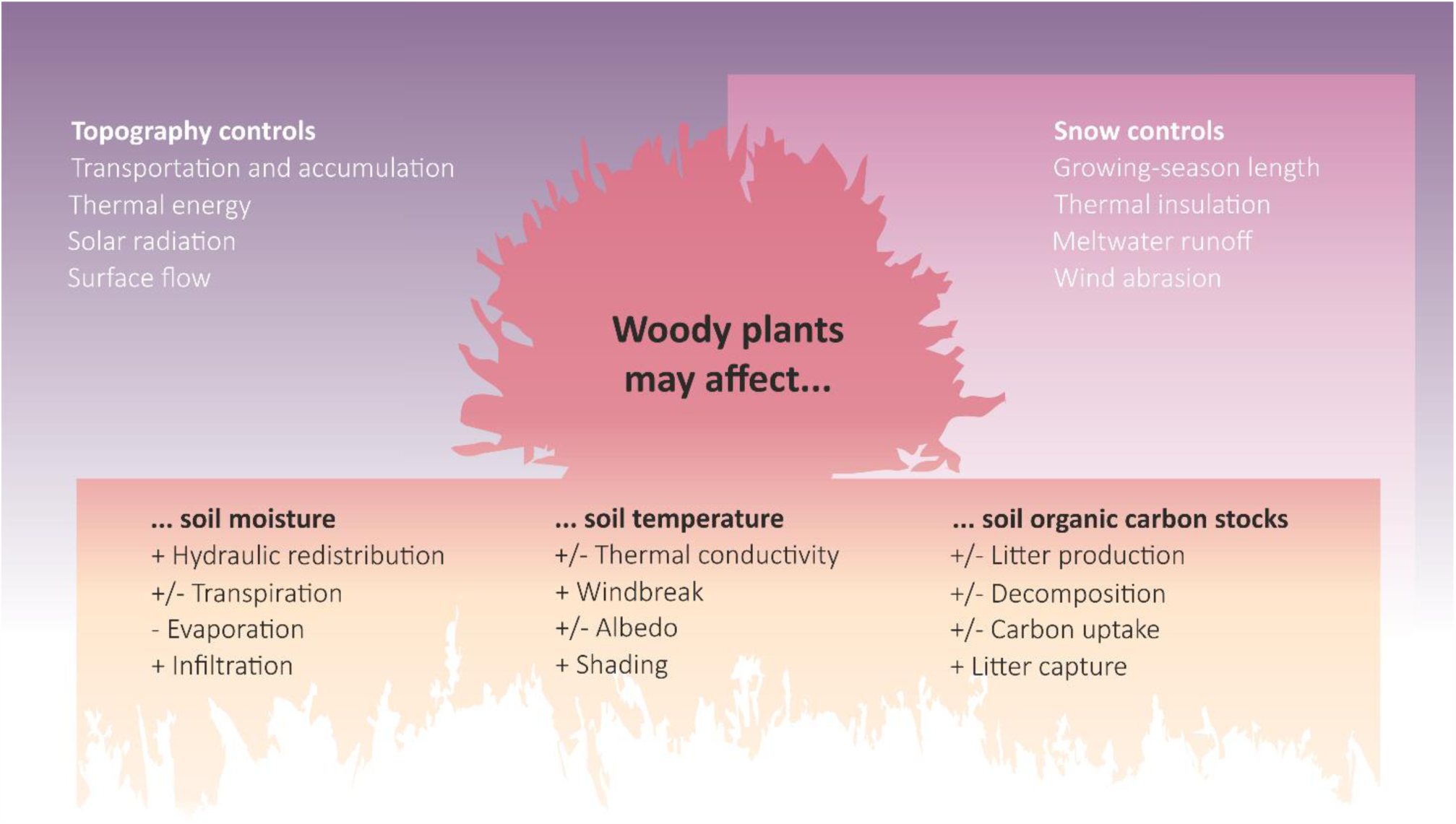
The mechanisms through which woody plants may affect soil conditions in tundra ecosystems.

Mountain tundra systems are characterised by a heterogeneous topography and seasonal snow cover (Billings, 1973). These factors play crucial roles on the spatial and temporal distribution of water and energy (Ayres et al., 2010; Kemppinen, Niittynen, Riihimaki, & Luoto, 2018; Williams, McNamara, & Chandler, 2009). Thus, winter conditions should be carefully considered in vegetation research conducted within seasonally snow-covered ecosystems (Niittynen & Luoto, 2018). Wintertime snow conditions may carry knock-on effects influencing summertime vegetation and soil properties by affecting the length of the growing season, nutrient availability, and the hydrothermal condition of soils (Semenchuk et al., 2015; Walker et al., 1999; Williams et al., 2009). Thus, topography and shrub–snow relations should be considered when investigating the impact of woody plants on tundra soil conditions (Figure 1) (Beck, Kalmbach, Joly, Stien, & Nilsen, 2005; Myers-Smith et al., 2011).

Soil hydrothermal conditions (i.e., the soil microclimate) have received increasing attention since they show spatio-temporal variability unrelated to macroclimatic trends (e.g., De Frenne et al., 2019; Gardner, Maclean, & Gaston, 2019; Lembrechts et al., 2019). Biological factors such as plants affect the hydrological and thermal properties of soils (Loranty et al., 2018; Robinson et al., 2019). For instance, vegetation causes drying in soils through transpiration, used for growth and reproduction (Horton & Hart, 1998), although water-use efficiency differs between plant species, functional groups, and ecosystems (Tang et al., 2014). In tundra, woody plants intercept rainfall, which, on a large scale, may decrease the overall water input into the ecosystem (Zwieback, Chang, Marsh, & Berg, 2019). However, vegetation can also cause opposing effects by providing shade, which in turn decreases not only the soil temperature, but also evaporation—that is, water vaporisation at the soil surface (Aalto, Scherrer, Lenoir, Guisan, & Luoto, 2018). Yet, woody plants in particular and their fundamental effects on the soil microclimate, especially on soil moisture, remain insufficiently investigated.

The soil and vegetation conditions are integral parts of the tundra carbon cycle (Cahoon et al., 2012). The slow decomposition rate of the cool tundra regions enables high-latitude ecosystems to store a large amount of carbon (Hugelius et al., 2014), roughly 50% of the global belowground organic carbon pool (Tarnocai et al., 2009). The expansion of woody plants is expected to increase the aboveground carbon storage within tundra ecosystems, which may further alter the rate of carbon cycling through changes in litter quality and biomass production (Myers-Smith & Hik, 2013). The shifts in vegetation composition towards a higher aboveground productivity may unexpectedly lead to decreasing belowground carbon storage, possibly decreasing the overall carbon storage of tundra ecosystems (Parker et al., 2015). Regardless of the direction of the effect, the expansion of woody plants is likely to affect the tundra carbon cycle carrying possibly consequences for the global carbon cycle (Cahoon et al., 2012). Yet, understanding how local soil organic carbon stocks respond to the expected changes in the expansion of woody plants remains woefully insufficient.

Dwarf shrub tundra forms a substantial part of the overall Arctic vegetation (Walker et al., 2017). As a shortcoming, investigations of the effects of woody plant expansion have primarily focused on tall, deciduous shrub species, whilst smaller, albeit more abundant, evergreen dwarf shrubs have received less attention (Vowles & Björk, 2018). Nonetheless, dwarf shrub expansion appears responsive to ongoing climate change (e.g. Buizer et al., 2012; Hallinger et al., 2010; Vuorinen et al., 2017). Here, we carried out an observational study in a dwarf shrub–dominated tundra and built a theory-based hierarchical model. We aimed to quantify the influence of woody plants on topsoil conditions (namely, soil moisture, soil temperature, and soil organic carbon stocks) after controlling for the effects of other factors (namely, the topography, wintertime snow depth, and overall vascular plant abundance). Here, we attempted to answer to the following overarching question: Do woody plants affect multiple soil conditions in tundra?

## Materials & methods

### Study area

We conducted this research in the Fennoscandian mountain tundra of north-western Finland (69°03’N 20°51’E), in a study area covering 3 km^2^ of a topographically heterogeneous landscape. The study design comprised 171 study plots (1 m^2^) located above the treeline between two mountains: Mount Saana and Mount Jehkas, which face north and south, respectively (Appendix 1). On average, July is the warmest (11.2°C) and wettest (73 mm) month at the study site according to measurements recorded at the Kilpisjärvi meteorological stations between 1981 and 2010 (Pirinen et al., 2012) (Appendix 2).

In Fennoscandia, the expansion of woody plants results from increasing summer temperatures and thicker snow cover during winter (Hallinger et al., 2010). In turn, the abundance and coverage of ground-dwelling dwarf shrubs have increased (Maliniemi, Kapfer, Saccone, Skog, & Virtanen, 2018; Wilson & Nilsson, 2009). The main vegetation type in our study area was dwarf shrub– dominated mountain heath with vegetation coverage consisting primarily of woody plant species. The dominant woody plant species are evergreen *Vaccinium vitis-idaea* and *Empetrum nigrum*, and deciduous *Betula nana* and *Vaccinium myrtillus*. Appendix 3 provides a full list of all woody plant species found in our study plots.

Topography plays an important role in the distribution of woody plants in tundra systems (Myers-Smith et al., 2011). The landscape in our study area is characterised by a rugged terrain, varying mesotopography, and predominantly thin soils (Bruun et al., 2006). The soil surface is covered by an organic soil layer, with a thickness varying by up to 70 cm (for more details, see Kemppinen et al., 2018). Appendix 1 provides maps describing the topography of the study area.

### Study design

In tundra, large areas of transition zones between habitats create variability in species composition and ecosystem functions (Fletcher et al., 2012). We selected the location of our study plots using a systematic grid approach covering the study area and its diversity of habitats as well as the transition zones between them (Appendix 1). We recorded the locations of the plots using a hand-held Global Navigation Satellite System receiver with an accuracy of up to ≤6 cm under optimal conditions (GeoExplorer GeoXH 6000 Series; Trimble Inc., Sunnyvale, CA, USA).

We excluded sites situated in river channels (with the exception of meltwater channels) or boulder fields, or those exposed to anthropogenic disturbance (i.e., trails). In addition to the systematic grid with a 50-m minimum distance between sites, we intentionally situated 25 sites in snow-bank environments and along windswept ridge tops to maximise the snow accumulation gradient in the data coverage. In all 171 plots, we recorded wintertime snow depth (at the maximum snow-depth timing in April), vegetation conditions (namely, plant coverage, woody plant dominance, and woody plant height), and the soil conditions (namely, soil moisture, soil temperature, and soil organic carbon stock).

### Soil data

#### Soil moisture

We measured the soil moisture during the 2017 growing season (following Kemppinen et al., 2018). We measured the soil moisture using a hand-held time-domain reflectometry sensor (FieldScout TDR 300; Spectrum Technologies Inc., USA). Moisture was measured up to a depth of 7.5 cm as the volumetric water content (VWC%). The resolution of the device is 0.1% with a reported accuracy of ±3.0 VWC%. We repeated the moisture measurements on five occasions (hereafter, campaigns; see Appendix 2). During each campaign, we took the mean over three measurements per plot to account for possible fine-scale moisture variation within a single plot. In subsequent analyses, we used the mean of these five campaigns to represent the overall soil moisture level during the growing season.

#### Soil temperature

We measured the soil temperature simultaneously with the soil moisture measurements. We used a hand-held digital temperature device (VWR-TD11; VWR International, USA). The resolution of the device is 0.1°C with a reported accuracy of ±0.8°C. Temperature was measured at a depth of up to 7.5 cm. We took the temperature measurement once from the centre of each plot, repeating this on five campaigns during the growing season (Appendix 2). In our analyses, we used the mean over the five campaigns to represent the soil temperature during the growing season. We corrected the possible effects of the timing of the measurements using data from miniature temperature loggers. Details regarding the soil temperature corrections appear in Appendix 4.

#### Soil organic carbon stock

We measured the depth of the organic soil layer from three points in each plot using a metal probe. In our analysis, we used the mean over the three measurement points per plot to represent the organic layer depth of each plot. We collected soil samples of approximately 1 dl of soil material from the organic and mineral soil layers using metal soil core cylinders (4–6 cm Ø, 5–7 cm in height). Samples from the organic soil layer were collected from the topsoil, with samples from the mineral soil layer taken from directly below the organic soil layer. The samples were collected from the same proximity of each plot between 1 and 31 August 2016 and 2017. A detailed description of the two sampling years appears in Appendix 5.

The laboratory analyses were carried out in the Laboratory of Geosciences and Geography and the Laboratory of Forest Sciences at the University of Helsinki. We used a freeze-drying method to remove soil moisture from the samples. The bulk density (kg/m^3^) was estimated by dividing the dry weight by the sample volume. The total carbon content (hereafter, C%) analyses were performed using elemental analysers (vario MICRO cube & vario MAX cube; Elementar Analysensysteme GmbH, Germany). Prior to the C% analysis, samples from the mineral soil layer were sieved through a 2-mm plastic sieve. Samples from the organic soil layer were homogenised by hammering the material into smaller particles.

Soil organic carbon stocks for the organic and mineral soil layers were estimated using Equation 1 (following Parker et al., 2015):

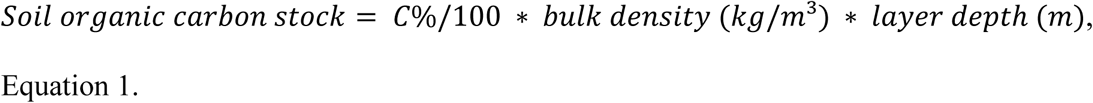

We used the median C% and bulk density of the samples from the mineral soil layer to estimate the soil organic carbon stocks in all mineral soil layers (3% for C%, n = 54, and 830 kg/m^3^ for bulk density, n = 70). We assumed that this would reliably allow us to determine the soil mineral layer stocks given the relatively low variability in C% (0.4–6.5%) and the bulk density (470–1400 kg/m^3^). We used the *in situ* measured mineral soil layer depth for each plot (relying on the mean over three measurements).

We did not measure C% from the samples taken from the organic soil layer from all sites. However, we measured the soil organic matter content (SOM%) using the loss-on-ignition method from the remaining sites (n = 52) (SFS3008, 1990). We calculated the soil organic matter stocks following Equation 1 and converted them into soil organic carbon stocks based on a relationship between C% and SOM% (Appendix 2). For this conversion, we used data from the study plots that had both C% and SOM% data based on samples from the organic soil layer (n = 118). The carbon fraction in the soil organic matter was calculated as 0.54 (R^2^ = 0.97), similar to Parker et al. (2015). This was then used to estimate the soil organic carbon stock in the organic layer. We arrived at this fraction by regressing the soil organic carbon stock in the organic soil layer using the soil organic matter stock in the organic soil layer without the intercept.

### Vegetation data

#### Woody plant dominance and height

We carried out a survey focusing on woody plants in which we visually estimated the coverage and measured the median height across three measurements of each woody plant species in all plots (Figure 2). Data on woody plants were collected from 23 to 27 July 2016 (146 plots) and completed between 16 and 19 July 2017 (25 plots). The relative coverage of the woody plants was calculated using the absolute coverage of all vascular plant species (hereafter, woody plant dominance).

**Figure 2.**
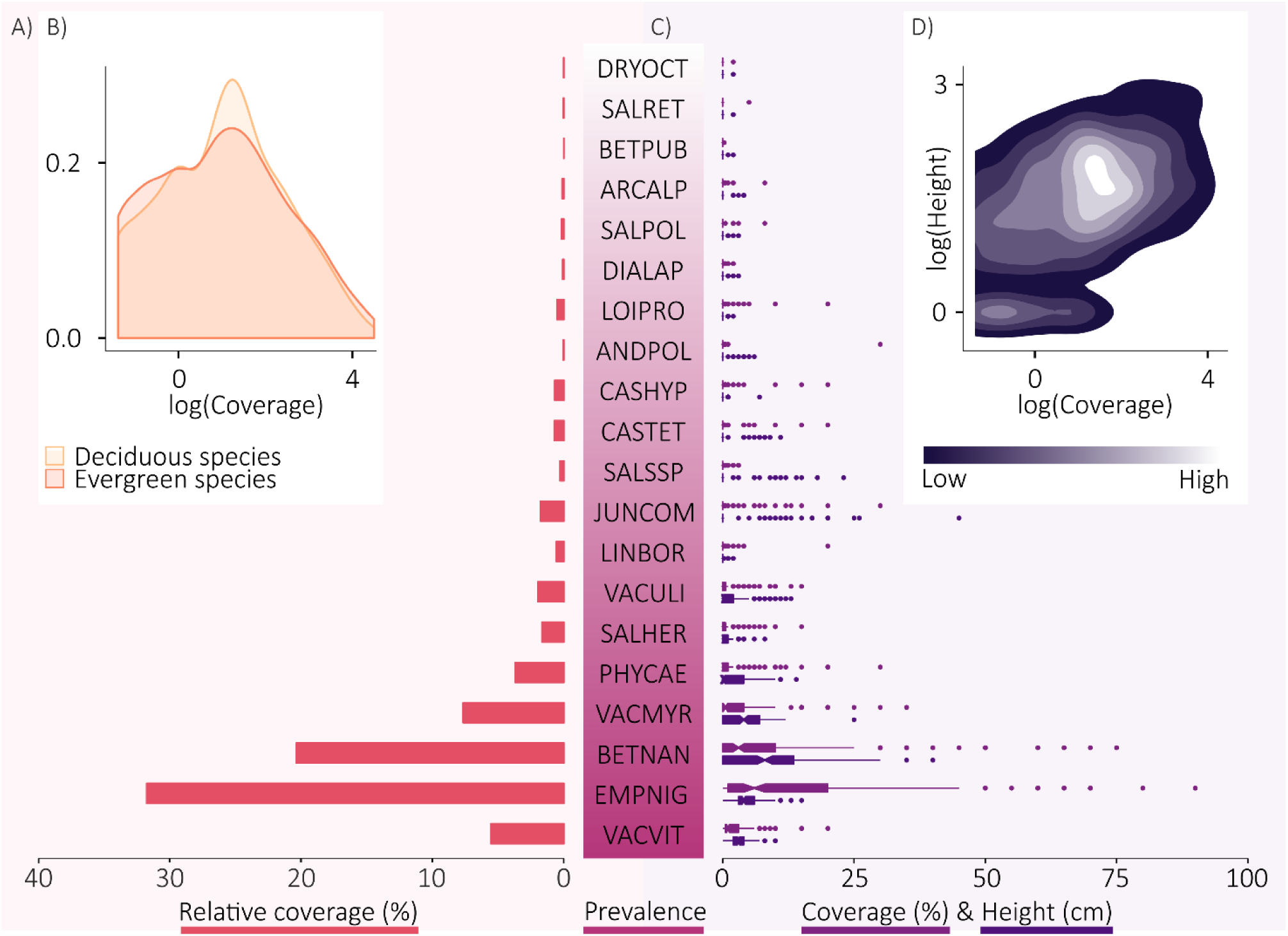
Woody plants of the dwarf shrub–dominated tundra. The relative coverage of the woody plants in relation to the total vascular plant coverage is shown as bars (A). The absolute woody plant coverage data divided between deciduous and evergreen species are presented as a frequency plot (B). The prevalence of the species ranks them from the most common (VACVIT = *Vaccinium vitis-idaea*) to the rarest (DRYOCT = *Dryas octopetala*) species occurring in the data. The hue intensity indicates the local point density. Species-specific coverage and median height data are shown as box plots (C). In the box plots, the notches and hinges represent the 25^th^, 50^th^, and 75^th^ percentiles. In addition, the whiskers represent the 95% percentile interval and the points represent the outliers. The variability of coverage and height data in relation to each other are shown in the density plot (D). Full species names appear in Appendix 3.

In total, we found 20 woody plant species (Figure 2; Appendix 3). In the height measurements, we excluded possible inflorescence to avoid any bias, particularly regarding the most ground-dwelling species such as *Cassiope hypnoides* and *Linnea borealis*. Besides the prostrate *Salix* species (namely, *Salix herbacea, S. polaris*, and *S. reticulate*), we grouped all erect *Salix* species into one category (*Salix* spp.) given the large amount of *Salix* hybridisation. We followed the taxonomy put forth by Hämet-Ahti, Suominen, Ulvinen, and Uotila (1998).

#### Plant coverage

The link between the vegetation and the soil conditions must be acknowledged, particularly in climate–soil–vegetation studies (Porporato & Rodriguez-Iturbe, 2002). In addition to the woody plants, a variety of graminoids and forbs may influence the soil conditions in tundra (Aalto et al., 2013). Thus, we included the effect of the overall vascular plant coverage in the model to separate out the effects of woody plant dominance from the effects of overall plant abundance as well as to distinguish the effects of woody plants on soil moisture from the effects that soil moisture have on vascular plants (Kemppinen, Niittynen, Aalto, le Roux, & Luoto, 2019).

Vascular plant coverage data were collected between 16 July and 8 August 2017. We carried out a vegetation survey, in which we visually estimated the species-specific percentage coverage of each vascular plant species in all plots. In the analysis, we use the plot-specific absolute coverage of all vascular plant species (hereafter, plant coverage). In densely vegetated plots, the plant coverage may exceed 100%, since vegetation may grow in several layers.

### Snow data

During winter, woody plants effectively capture drifting snow, which may alter the distribution of snow (Lafleur & Humphreys, 2018). Consequently, if trapped, the accumulation of snow may lead to thicker and more insulative snow cover in and around shrub patches compared to shrubless patches (Sturm et al., 2001). Yet, this is unlikely for dwarf shrubs due to their prostrate stature (Vowles & Björk, 2018). Nonetheless, snow cover—or the lack of it—may have consequences for all woody plants regardless of their size (Hallinger et al., 2010; Phoenix & Bjerke, 2016).

We measured the snow depth to represent the winter conditions in our study system (Aalto et al., 2018). We collected snow depth data between 13 and 17 April 2017 during the maximum snow depth during the season. We measured the snow depth from the centre of each plot using an aluminium probe.

### Topographic data

We obtained the LiDAR-based digital terrain model (DTM) at a 2-m resolution from the open file service of the National Land Survey of Finland (NLS, 2019). Based on the 2-m resolution DTM, we calculated the elevation, radiation, the topographic wetness index (TWI), and the topographic position index (TPI) (Appendix 1).

#### Elevation

Elevation represents the macroclimatic conditions of the system, which we assumed were reflected in the snow depth, vegetation, and soil variables for our study system. Differences in the elevation are followed by several environmental gradients, such as air temperature due to a lapse rate (i.e., general decrease in the air temperature with altitude). This, in turn, affects the coverage and height of the vegetation (Riihimäki, Heiskanen, & Luoto, 2017), as well as the soil formation, soil microclimate, and microbial and cryogenic activity (Amundson, Chadwick, & Sowers, 1989; Dahlgren, Boettinger, Huntington, & Amundson, 1997; Trumbore, Chadwick, & Amundson, 1996).

#### Radiation

Solar radiation is distributed unevenly across the landscape due to the topographic variability. This creates a strong radiation gradient between the north- and south-facing slopes in the study area (le Roux, Aalto, & Luoto, 2013). We assumed that solar radiation positively correlated with both the coverage and height of the vegetation variables as well as with the soil temperature (Aalto, le Roux, & Luoto, 2014; Winkler et al., 2016).

We calculated the potential incoming solar radiation based on the three months of the growing season (June, July, and August) using a tool in SAGA GIS v. 2.3.2 that measures the potential incoming solar radiation. We considered the possible shadow effect of obstructing topographic features using the sky view–factor option (Böhner & Antonic, 2009). We calculated the position of the Sun for every fifth day using a 4-h interval. We used the lumped atmosphere option to calculate the atmospheric transmittance.

#### Topographic wetness index

The topography controls the surface water flow and the accumulation of it (Beven & Kirkby, 1979). We assumed that the water flow and accumulation for the landscape positively correlated with both the coverage and the height of the vegetation, as well as the soil moisture and soil organic carbon stock (Kemppinen et al., 2018; le Roux et al., 2013).

We calculated the topographic wetness index [TWI = ln (SCA / local slope)], a proxy for the water flow and accumulation, which can be used to model the variation in the soil moisture at fine-spatial scales (Kemppinen et al., 2018). The total catchment area (TCA) was calculated from a filled DTM using the multiple flow–direction algorithm (Freeman, 1991; Wang & Liu, 2006), which performs best in predicting volumetric water content in mountain tundra (Riihimäki, Kemppinen, Kopecký, & Luoto, 2019). The specific catchment area (SCA) was calculated assuming that the flow width equals the grid resolution (2 m) (i.e., SCA = TCA/2). The local slope was calculated using the algorithm by Zevenbergen and Thorne (1987).

#### Topographic position index

Topographic position affects the erosion, transportation, and accumulation of matter and energy in the landscape. We assumed that the topographic position associates with snow depth specifically in the study system (Billings & Bliss, 1959).

We calculated the topographic position index (TPI), based on the elevation difference between a plot and the surrounding elevation along a given radius (Agren, Lidberg, Stromgren, Ogilvie, & Arp, 2014). This describes the position of the plot on a topographic gradient—that is, a plot located on a ridge top (positive values), in a depression (negative), or on a slope or flat ground (close to zero). We calculated TPI using 30-m radii based on an unfilled DTM (following Kemppinen et al., 2018).

### Statistical analyses

We used structural equation modelling (SEM) to understand the hierarchy of the environmental variables controlling the soil conditions in tundra systems. SEM is a statistical modelling method that can be used to understand complex multivariate relationships, such as indirect effects and cascading effects (Grace, Anderson, Olff, & Scheiner, 2010). SEM produces a single causal network in which several response variables and predictors are combined by probabilistic models (Lefcheck, 2016). This enables the testing of hypothesised causal relationships, here presented as arrows. In SEM, a variable can be both a predictor and a response variable. This enables determining if a predictor has a direct or mediating effect—that is, if a variable serves as a mediator. Thus, SEM allows the simultaneous evaluation of several causal structures. Results are expressed as standardised regression coefficients, which allows for the direct comparison of the estimated effects (Lefcheck, 2016). We used the *piecewiseSEM* package in R, version 3.5.1 (Lefcheck, 2016; R Development Core Team, 2016).

We log-transformed all three vegetation variables (plant coverage, woody plant dominance, and woody plant height) as well as the soil moisture and soil organic carbon stock variables to reduce the heteroscedasticity in the component models. This decision was based on the visual interpretation of the histograms of the linear model residuals for each response variable.

We also calculated the bivariate Spearman correlations between all variables. Due to the high correlation (−0.72) between the snow depth and TPI variables (Appendix 6), we excluded TPI from those models that used snow depth as a predictor (i.e., all vegetation and soil models).

We fitted all hypothesised paths (i.e., component models) using a first-order multiple linear regression model. The paths included both the direct pathways from the predictors (topography) and pathways through the mediators (snow depth and vegetation variables) to the response variables (soil variables). We created SEM consisting of seven component models as follows. 1) Snow depth was modelled using elevation and TPI. 2) Plant coverage, 3) woody plant dominance, and 4) woody plant height were modelled using elevation, radiation, TWI, and snow depth. 5) Soil moisture, 6) soil temperature, and 7) soil organic carbon stock were modelled using all predictors and mediators (for a detailed model structure, see Appendix 7).

We assumed that correlations might exist between the error terms of the soil and vegetation variables. Thus, we updated the model to account for all possible residual correlations between the vegetation variables and between the soil variables.

### Data deposition

Field data will be made openly available in Zenodo once this manuscript is accepted, and we will include the link here.

## Results

The R^2^ values for the seven component models ranged from 0.09 to 0.54 (Figure 3; for detailed results of the model, see Appendix 7). According to tests of directed separation, we excluded from the model only one path, which was a link between woody plant height and TPI.

**Figure 3.**
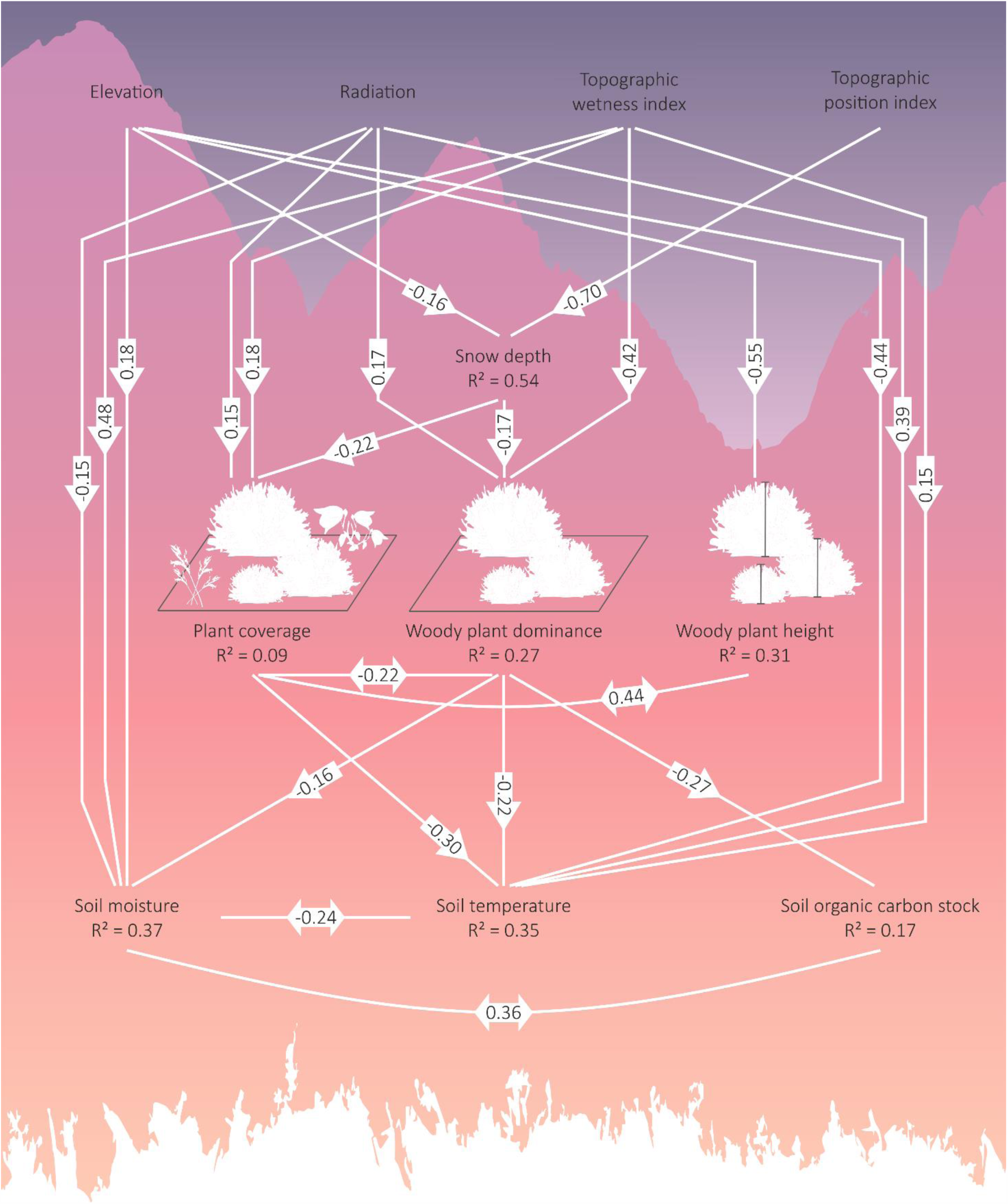
Woody plants constructing tundra soils. Results of a structural equation model with seven component models. Here, we only present relationships with *P* < 0.05. The arrows show the standardised regression slopes associated with the links. Double-headed arrows represent correlations between error terms. For further details and relationships with *P* ≥ 0.05, see Appendix 7.

Soil temperature was the only soil variable that linked to plant coverage (standardised coefficient = −0.30, *P* = 0.00). Woody plant dominance negatively correlated with all soil conditions: soil moisture, soil temperature, and soil organic carbon stock (standardised coefficients = −0.16; −0.22; - 0.27, *P* = 0.04; 0.00; 0.00). None of the soil conditions were affected by woody plant height. The covariance between soil moisture and soil organic carbon stock was positive.

We found no significant direct link (*P* < 0.05) for snow depth to the soil conditions. Yet, the snow depth negatively affected plant coverage and woody plant dominance (standardised coefficients = - 0.22; −0.17, *P* = 0.01; 0.02), respectively.

## Discussion

We found that woody plants such as dwarf shrubs affect the soil moisture, soil temperature, and soil carbon stock distribution in the tundra ecosystem. In addition to woody plants, we considered the direct and indirect effects of the topography, wintertime snow depth, and overall vascular plant coverage on the tundra soil conditions.

### Effects of woody plants on the soil moisture

We found that the dominance of woody plants directly related to less soil moisture in the topsoil. However, our results indicated no relationship between woody plant height and soil moisture. This suggests that the expansion of woody plants may carry different consequences on the soil moisture depending upon whether the expansion is realised through the coverage or the height of woody plants.

Our findings are relevant in light of previous studies, which found that the expansion of woody plants may lead to an increased evapotranspiration and intercepting precipitation (Bonfils et al., 2012; Pearson et al., 2013; Zwieback et al., 2019). Previous studies reached conclusions similar to our fine-scale investigation: that is, soil moisture is generally lower in woody plant habitats compared to other tundra habitats (Ge, Lafleur, & Humphreys, 2017; Lafleur & Humphreys, 2018).

The negative effect of woody plants on the soil moisture could be explained by an increased transpiration. Transpiration also appears sensitive to the height of woody plants, since taller plants transpire more than smaller plants (Bonfils et al., 2012). In tundra, evergreen woody plants may cause less evaporation, since they cast shade on the soil beneath them throughout the growing season. Yet, deciduous woody plant transpiration is affected by phenology, as transpiration may overcome evaporation as soon as buds burst into leaves (Bonfils et al., 2012). The expansion of woody plants can increase water vapour in the atmosphere, because woody plants transpire water more efficiently than barren tundra (Swann, Fung, Levis, Bonan, & Doney, 2010). An increased transpiration, in turn, may amplify the greenhouse effect of woody plants alongside changes in the albedo (Swann et al., 2010).

We found no relationship between vascular plant coverage and soil moisture. This shows that we could 1) separate the effects of woody plants from the overall effects of vascular plants and 2) distinguish between the effects of woody plants on the soil moisture from the effects of soil moisture on vascular plants. Thus, the expansion of woody plants may affect the soil moisture beyond that resulting from higher overall vegetation coverage. Therefore, we stress that it is crucial to consider carefully which plant–soil feedbacks are essential in next-generation soil hydrology models (Porporato & Rodriguez-Iturbe, 2002; Robinson et al., 2019).

### Effects of woody plants on soil temperature

We found that plants in general lower the soil temperatures in tundra, given that both the overall plant coverage and woody plant dominance decreased the soil temperature. In treeless ecosystems (such as tundra), vegetation in general cools the soils below through shading (Aalto et al., 2013; Myers-Smith & Hik, 2013). In addition, both deciduous and evergreen woody plants appear to carry a cooling effect on soil temperatures during the growing season (Loranty et al., 2018).

The interactions between the macroclimate, vegetation, and microclimate are strongly linked to each other. Because the macroclimate in tundra regions is warming, woody plants are expected to expand (Myers-Smith et al., 2015). In turn, they alter the microclimate as our results indicate. Woody plants can carry strong effects on the radiation and the mixing of air; thus, abundant vegetation may create heterogeneous microclimates (De Frenne et al., 2019). Tall, erect species can potentially cause wind turbulence, and, in turn, enable the release of latent heat from the soil to the open atmosphere (Bonfils et al., 2012). Small, prostrate species such as cushion plants, however, grow closer to ground and are aerodynamic compared to taller species, which may lead to less turbulence at the ground surface.

### Effects of woody plants on soil organic carbon stock

We found that the dominance of woody plants directly relates to lower soil organic carbon stocks. Yet, we found no relationship between woody plant height and the soil organic carbon stock (Figure 3), although we did detect a significant positive bivariate correlation between these two variables (Appendix 6). These findings show that the expansion of woody plants (i.e., increasing coverage or height) may influence the tundra carbon cycle in various ways. This may explain why no consensus currently exists regarding how the expansion of woody plants may affect the tundra carbon cycle.

Other studies found that the expansion of woody plants may lead to either an increase (Qian, Joseph, & Zeng, 2010) or decrease (Cahoon et al., 2012) in the soil organic carbon, or carry no effect at all (Sistla et al., 2013). Woody plants can cycle carbon more rapidly compared to herbaceous plants (Cahoon et al., 2012; Lafleur & Humphreys, 2018). The expansion of woody plants in tundra may result in larger, yet less decomposable, litter inputs into the soil (DeMarco, Mack, & Bret-Harte, 2014). By contrast, in warmer macroclimatic conditions, large-scale expansion may lead to faster soil decomposition depending upon the type of habitat the woody plant species expands to (Ge et al., 2017). Plant root properties and rhizosphere processes also influence soil organic carbon stocks (Parker et al., 2015). For example, root longevity (i.e., age) is lower in graminoids than in shrubs (Iversen et al., 2015), resulting in slower carbon cycling in soils under shrub-dominated vegetation. In addition, the roots of woody plants are shallower than those of graminoids; thus, an increase in woody plant coverage may result in smaller carbon inputs from roots to deeper soil layers (Ylänne, Olofsson, Oksanen, & Stark, 2018).

We speculate that our findings can be explained by a combination of all of these factors. Woody plant dominance may lead to decreased soil organic carbon stocks due to a faster decomposition, a more active soil microbiota, and smaller carbon inputs from roots to the soil (Ge et al., 2017). Our results might also be explained by the covariance of soil organic carbon stocks with the soil moisture. In this study, the wettest soils were most often dominated by non-woody plants. Specifically, the plots with tall woody plants had a relatively low coverage, possibly explaining why we found no link between woody plant height and soil organic carbon stock in the model.

Quantifying soil organic carbon may introduce some uncertainties to the model (Hugelius et al., 2014). Some of our study sites were located in tundra meadows (<10%), in which the uppermost part (roughly 0–30 cm) of the soil can in some parts represent a mixture of both organic and mineral layers. Thus, distinguishing the organic layer from the mineral might be reflected in the estimations of the soil organic carbon stocks.

### Indirect effects of snow on soil conditions

We found that wintertime snow depth affected summertime soil conditions indirectly through the vegetation properties. These results show that snow depth directly correlated with overall plant coverage and woody plant dominance. Yet, we found no direct effects of snow depth on the soil conditions.

In high-latitude mountainous landscapes, the winter conditions have far-reaching effects long into the summer, since a thick snow cover affects the vegetation through plant development and productivity (Billings & Bliss, 1959), and by affecting plant functional traits, such as the specific leaf area or the dry-matter content (Happonen et al., 2019). Wintertime shrub–snow relationships affect wintertime soil conditions, such as wintertime soil temperatures, shown to positively feedback on shrubs (Myers-Smith & Hik, 2013; Sturm et al., 2005). For example, warming winter conditions are as relevant as summertime changes in the temperature for the growth of *Betula nana* (Hollesen et al., 2015).

Our results regarding snow were unexpected, since previous studies found that wintertime snow conditions affect multiple soil conditions, particularly the spatio-temporal variability of the growing-season soil moisture (Ayres et al., 2010), even when controlling for the effects of topographical variation (Williams et al., 2009). Nevertheless, a careful consideration of snow in plant–soil interactions remains crucial wherever snow is present (Niittynen, Heikkinen, & Luoto, 2018). In fact, several experimental and observational studies have linked wintertime snow patterns to ecosystem processes through changing the soil moisture availability (e.g. Ayres et al., 2010; Barichivich et al., 2014; Plaza et al., 2019).

### In summary

We investigated the effects of woody plants on multiple soil conditions in tundra. Through this hierarchical approach, we controlled for the influence of other possible factors—namely, the topography, wintertime snow depth, and overall vascular plant conditions. Our study benefitted from a hierarchical framework, since previous studies found it challenging to test whether the effects found resulted from changing vegetation properties or from, for instance, the underlying topography (Crofts, Drury, & McLaren, 2018). Thus, we recommend future studies consider SEM in their analyses when investigating hierarchical systems (Lefcheck, 2016).

Our results suggest that the dominance of woody plants negatively affects the soil moisture, soil temperature, and soil organic carbon stocks. Yet, the height of woody plants did not affect the soil conditions in our model when considering the coverage of woody plants. These findings indicate that the effects of woody plants on tundra soil conditions depend upon whether expansion occurs in terms of the coverage or height. Thus, future studies should consider quantifying multiple dimensions of woody plants to describe their expansion in tundra ecosystems.

## Acknowledgements

The authors thank all past and present members of the BioGeoClimate Modelling Lab and the Arctic Microbial Ecology group at the University of Helsinki for their assistance during field and laboratory work, and the staff at the Kilpisjärvi Biological Station for their support. Dr. Aleksandra M. Lewandowska and Dr. Dorothee Hodapp are greatly acknowledged for their help and encouragement regarding SEM. We are also grateful to Vanessa Fuller for assistance with English-language revision. We also thank Professor Gustaf Olsson for sharing his knowledge of scientific writing. The National Land Survey of Finland and the Finnish Meteorological Institute gratefully provided the LiDAR and meteorological data.

## Permissions

Permission to carry out fieldwork was granted by Metsähallitus.

## References

Aalto, J., le Roux, P. C., & Luoto, M. (2013). Vegetation Mediates Soil Temperature and Moisture in Arctic-Alpine Environments. Arctic Antarctic and Alpine Research, 45(4), 429–439. doi:10.1657/1938-4246-45.4.429

Aalto, J., le Roux, P. C., & Luoto, M. (2014). The meso-scale drivers of temperature extremes in high-latitude Fennoscandia. Climate Dynamics, 42(1-2), 237–252. doi:10.1007/s00382-012-1590-y

Aalto, J., Scherrer, D., Lenoir, J., Guisan, A., & Luoto, M. (2018). Biogeophysical controls on soil-atmosphere thermal differences: implications on warming Arctic ecosystems. Environmental Research Letters, 13(7), 074003. doi:ARTN07400310.1088/1748-9326/aac83e

Agren, A. M., Lidberg, W., Stromgren, M., Ogilvie, J., & Arp, P. A. (2014). Evaluating digital terrain indices for soil wetness mapping - a Swedish case study. Hydrology and Earth System Sciences, 18(9), 3623–3634. doi:10.5194/hess-18-3623-2014

Amundson, R. G., Chadwick, O. A., & Sowers, J. M. (1989). A comparison of soil climate and biological activity along an elevation gradient in the eastern Mojave Desert. Oecologia, 80(3), 395–400. doi:10.1007/BF00379042

Ayres, E., Nkem, J. N., Wall, D. H., Adams, B. J., Barrett, J. E., Simmons, B. L., … Fountain, A. G. (2010). Experimentally increased snow accumulation alters soil moisture and animal community structure in a polar desert. Polar Biology, 33(7), 897–907. doi:10.1007/s00300-010-0766-3

Barichivich, J., Briffa, K. R., Myneni, R., van der Schrier, G., Dorigo, W., Tucker, C. J., … Melvin, T. M. (2014). Temperature and Snow-Mediated Moisture Controls of Summer Photosynthetic Activity in Northern Terrestrial Ecosystems between 1982 and 2011. Remote Sensing, 6(2), 1390–1431. doi:10.3390/rs6021390

Beck, P. S. A., Kalmbach, E., Joly, D., Stien, A., & Nilsen, L. (2005). Modelling local distribution of an Arctic dwarf shrub indicates an important role for remote sensing of snow cover. Remote Sensing of Environment, 98(1), 110–121. doi:10.1016/j.rse.2005.07.002

Beven, K. J., & Kirkby, M. J. (1979). A physically based, variable contributing area model of basin hydrology. Hydrological Sciences Bulletin, 24:1, 43–69.

Billings, W. D. (1973). Arctic and Alpine Vegetations - Similarities, Differences, and Susceptibility to Disturbance. Bioscience, 23(12), 697–704. doi:Doi 10.2307/1296827

Billings, W. D., & Bliss, L. C. (1959). An Alpine Snowbank Environment and Its Effects on Vegetation, Plant Development, and Productivity. Ecology, 40(3), 388–397. doi:Doi 10.2307/1929755

Bonfils, C. J. W., Phillips, T. J., Lawrence, D. M., Cameron-Smith, P., Riley, W. J., & Subin, Z. M. (2012). On the influence of shrub height and expansion on northern high latitude climate. Environmental Research Letters, 7(1), 015503. doi:Artn 015503 10.1088/1748-9326/7/1/015503

Bruun, H. H., Moen, J., Virtanen, R., Grytnes, J. A., Oksanen, L., & Angerbjorn, A. (2006). Effects of altitude and topography on species richness of vascular plants, bryophytes and lichens in alpine communities. Journal of Vegetation Science, 17(1), 37–46. doi:DOI 10.1111/j.1654-1103.2006.tb02421.x

Buizer, B., Weijers, S., van Bodegom, P. M., Alsos, I. G., Eidesen, P. B., van Breda, J., … Rozema, J. (2012). Range shifts and global warming: ecological responses of Empetrum nigrum L. to experimental warming at its northern (high Arctic) and southern (Atlantic) geographical range margin. Environmental Research Letters, 7(2), 025501. doi:Artn 025501 10.1088/1748-9326/7/2/025501

Böhner, J., & Antonic, O. (2009). Land surface parameters specific to topo-climatology. Amsterdam: Elsevier.

Cahoon, S. M., Sullivan, P. F., Shaver, G. R., Welker, J. M., Post, E., & Holyoak, M. (2012). Interactions among shrub cover and the soil microclimate may determine future Arctic carbon budgets. Ecology Letters, 15(12), 1415–1422. doi:10.1111/j.1461-0248.2012.01865.x

Carrer, M., Pellizzari, E., Prendin, A. L., Pividori, M., & Brunetti, M. (2019). Winter precipitation - not summer temperature - is still the main driver for Alpine shrub growth. Science of the Total Environment, 682, 171–179. doi:10.1016/j.scitotenv.2019.05.152

Chapin, F. S., 3rd, Sturm, M., Serreze, M. C., McFadden, J. P., Key, J. R., Lloyd, A. H., … Welker, J. M. (2005). Role of land-surface changes in arctic summer warming. Science, 310(5748), 657–660. doi:10.1126/science.1117368

Crofts, A. L., Drury, D. O., & McLaren, J. R. (2018). Changes in the understory plant community and ecosystem properties along a shrub density gradient. Arctic Science, 4(4), 485–498. doi:10.1139/as-2017-0026

Dahlgren, R. A., Boettinger, J. L., Huntington, G. L., & Amundson, R. G. (1997). Soil development along an elevational transect in the western Sierra Nevada, California. Geoderma, 78 (3-4), 207–236. doi:Doi 10.1016/S0016-7061(97)00034-7

De Frenne, P., Zellweger, F., Rodriguez-Sanchez, F., Scheffers, B. R., Hylander, K., Luoto, M., … Lenoir, J. (2019). Global buffering of temperatures under forest canopies. Nat Ecol Evol, 3(5), 744–749. doi:10.1038/s41559-019-0842-1

DeMarco, J., Mack, M. C., & Bret-Harte, M. S. (2014). Effects of arctic shrub expansion on biophysical vs. biogeochemical drivers of litter decomposition. Ecology, 95(7), 1861–1875.

Fletcher, B. J., Gornall, J. L., Poyatos, R., Press, M. C., Stoy, P. C., Huntley, B., … Phoenix, G. K. (2012). Photosynthesis and productivity in heterogeneous arctic tundra: consequences for ecosystem function of mixing vegetation types at stand edges. Journal of Ecology, 100(2), 441–451. doi:10.1111/j.1365-2745.2011.01913.x

Freeman, T. G. (1991). Calculating Catchment-Area with Divergent Flow Based on a Regular Grid. Computers & Geosciences, 17(3), 413–422. doi:Doi 10.1016/0098-3004(91)90048-I

Gardner, A. S., Maclean, I. M., & Gaston, K. J. (2019). Climatic predictors of species distributions neglect biophysiologically meaningful variables. Diversity & Distributions.

Ge, L., Lafleur, P. M., & Humphreys, E. R. (2017). Respiration from soil and ground cover vegetation under tundra shrubs. Arctic Antarctic and Alpine Research, 49(4), 537–550. doi:10.1657/Aaar0016-064

Grace, J. B., Anderson, T. M., Olff, H., & Scheiner, S. M. (2010). On the specification of structural equation models for ecological systems. Ecological Monographs, 80(1), 67–87. doi:Doi 10.1890/09-0464.1

Hallinger, M., Manthey, M., & Wilmking, M. (2010). Establishing a missing link: warm summers and winter snow cover promote shrub expansion into alpine tundra in Scandinavia. New Phytol, 186(4), 890–899. doi:10.1111/j.1469-8137.2010.03223.x

Happonen, K., Aalto, J., Kemppinen, J., Niittynen, P., Virkkala, A.-M., & Luoto, M. J. B. (2019). Snow is an important control of plant community functional composition. 564583.

Hollesen, J., Buchwal, A., Rachlewicz, G., Hansen, B. U., Hansen, M. O., Stecher, O., & Elberling, B. J. G. C. B. (2015). Winter warming as an important co-driver for Betula nana growth in western Greenland during the past century. 21(6), 2410–2423.

Horton, J. L., & Hart, S. C. (1998). Hydraulic lift: a potentially important ecosystem process. Trends Ecol Evol, 13(6), 232–235.

Hugelius, G., Strauss, J., Zubrzycki, S., Harden, J. W., Schuur, E. A. G., Ping, C. L., … Kuhry, P. (2014). Estimated stocks of circumpolar permafrost carbon with quantified uncertainty ranges and identified data gaps. Biogeosciences, 11(23), 6573–6593. doi:10.5194/bg-11-6573-2014

Hämet-Ahti, L., Suominen, J., Ulvinen, T., & Uotila, P. (1998). Retkeilykasvio (Field Flora of Finland, in Finnish). Finnish Museum of Natural History, Botanical Museum. 656p.

Iversen, C. M., Sloan, V. L., Sullivan, P. F., Euskirchen, E. S., McGuire, A. D., Norby, R. J., … Wullschleger, S. D. (2015). The unseen iceberg: plant roots in arctic tundra. New Phytol, 205(1), 34–58. doi:10.1111/nph.13003

Jeffers, E. S., Bonsall, M. B., Watson, J. E., & Willis, K. J. (2012). Climate change impacts on ecosystem functioning: evidence from an Empetrum heathland. New Phytol, 193(1), 150–164. doi:10.1111/j.1469-8137.2011.03907.x

Kemppinen, J., Niittynen, P., Aalto, J., le Roux, P. C., & Luoto, M. (2019). Water as a resource, stress and disturbance shaping tundra vegetation. Oikos, Accepted Author Manuscript. doi:10.1111/oik.05764

Kemppinen, J., Niittynen, P., Riihimaki, H., & Luoto, M. (2018). Modelling soil moisture in a high-latitude landscape using LiDAR and soil data. Earth Surface Processes and Landforms, 43(5), 1019–1031. doi:10.1002/esp.4301

Lafleur, P. M., & Humphreys, E. R. (2018). Tundra shrub effects on growing season energy and carbon dioxide exchange. Environmental Research Letters, 13(5), 055001. doi:ARTN 055001 10.1088/1748-9326/aab863

le Roux, P. C., Aalto, J., & Luoto, M. (2013). Soil moisture’s underestimated role in climate change impact modelling in low-energy systems. Glob Chang Biol, 19(10), 2965–2975. doi:10.1111/gcb.12286

Lefcheck, J. S. (2016). PIECEWISESEM: Piecewise structural equation modelling in R for ecology, evolution, and systematics. Methods in Ecology and Evolution, 7(5), 573–579. doi:10.1111/2041-210x.12512

Lembrechts, J. J., Lenoir, J., Roth, N., Hattab, T., Milbau, A., Haider, S., … Biogeography. (2019). Comparing temperature data sources for use in species distribution models: From in-situ logging to remote sensing.

Loranty, M. M., Abbott, B. W., Blok, D., Douglas, T. A., Epstein, H. E., Forbes, B. C., … Malhotra, A. (2018). Reviews and syntheses: Changing ecosystem influences on soil thermal regimes in northern high-latitude permafrost regions (1810-6285). Retrieved from

Maliniemi, T., Kapfer, J., Saccone, P., Skog, A., & Virtanen, R. (2018). Long-term vegetation changes of treeless heath communities in northern Fennoscandia: Links to climate change trends and reindeer grazing. Journal of Vegetation Science.

Mod, H. K., & Luoto, M. (2016). Arctic shrubification mediates the impacts of warming climate on changes to tundra vegetation. Environmental Research Letters, 11(12), 124028. doi:Artn 124028 10.1088/1748-9326/11/12/124028

Myers-Smith, I. H., Elmendorf, S. C., Beck, P. S. A., Wilmking, M., Hallinger, M., Blok, D., … Vellend, M. (2015). Climate sensitivity of shrub growth across the tundra biome. Nature Climate Change, 5(9), 887–891. doi:10.1038/nclimate2697

Myers-Smith, I. H., Forbes, B. C., Wilmking, M., Hallinger, M., Lantz, T., Blok, D., … Hik, D. S. (2011). Shrub expansion in tundra ecosystems: dynamics, impacts and research priorities. Environmental Research Letters, 6(4), 15. doi:Artn 045509 10.1088/1748-9326/6/4/045509

Myers-Smith, I. H., & Hik, D. S. (2013). Shrub canopies influence soil temperatures but not nutrient dynamics: An experimental test of tundra snow-shrub interactions. Ecol Evol, 3(11), 3683–3700. doi:10.1002/ece3.710

Niittynen, P., Heikkinen, R. K., & Luoto, M. (2018). Snow cover is a neglected driver of Arctic biodiversity loss. Nature Climate Change, 8(11), 997-+. doi:10.1038/s41558-018-0311-x

Niittynen, P., & Luoto, M. (2018). The importance of snow in species distribution models of arctic vegetation. Ecography, 41(6), 1024–1037. doi:10.1111/ecog.03348

NLS. (2019). National Land Survey of Finland. https://www.maanmittauslaitos.fi/en/maps-and-spatial-data/expert-users/product-descriptions/elevation-model-2-m, Access date: 19.4.2019.

Parker, T. C., Subke, J. A., & Wookey, P. A. (2015). Rapid carbon turnover beneath shrub and tree vegetation is associated with low soil carbon stocks at a subarctic treeline. Global Change Biology, 21(5), 2070–2081. doi:10.1111/gcb.12793

Pearson, R. G., Phillips, S. J., Loranty, M. M., Beck, P. S. A., Damoulas, T., Knight, S. J., & Goetz, S. J. (2013). Shifts in Arctic vegetation and associated feedbacks under climate change. Nature Climate Change, 3(7), 673–677. doi:10.1038/Nclimate1858

Phoenix, G. K., & Bjerke, J. W. (2016). Arctic browning: extreme events and trends reversing arctic greening. Glob Chang Biol, 22(9), 2960–2962. doi:10.1111/gcb.13261

Pirinen, P., Simola, H., Aalto, J., Kaukoranta, J.-P., Karlsson, P., & Ruuhela, R. (2012). Climatological statistics of Finland 1981–2010. Retrieved from Helsinki:

Plaza, C., Pegoraro, E., Bracho, R., Celis, G., Crummer, K. G., Hutchings, J. A., … Salmon, V. G. (2019). Direct observation of permafrost degradation and rapid soil carbon loss in tundra. Nature Geoscience, 12(8), 627.

Porporato, A., & Rodriguez-Iturbe, I. (2002). Ecohydrology - a challenging multidisciplinary research perspective. Hydrological Sciences Journal-Journal Des Sciences Hydrologiques, 47(5), 811–821. doi:Doi 10.1080/02626660209492985

Qian, H. F., Joseph, R., & Zeng, N. (2010). Enhanced terrestrial carbon uptake in the Northern High Latitudes in the 21st century from the Coupled Carbon Cycle Climate Model Intercomparison Project model projections. Global Change Biology, 16(2), 641–656. doi:10.1111/j.1365-2486.2009.01989.x

R Development Core Team. (2016). The R Project for Statistical Computing, Vienna, Austria. R Development Core Team. Retrieved from <https://www.r-project.org/>

Riihimäki, H., Heiskanen, J., & Luoto, M. (2017). The effect of topography on arctic-alpine aboveground biomass and NDVI patterns. International journal of applied earth observation geoinformation, 56, 44–53.

Riihimäki, H., Kemppinen, J., Kopecký, M., & Luoto, M. (2019). Modelling performance of the Topographic Wetness Index is affected by grid resolution and flow-routing algorithm. Under review.

Robinson, D. A., Hopmans, J. W., Filipovic, V., van der Ploeg, M., Lebron, I., Jones, S. B., … Tuller, M. (2019). Global environmental changes impact soil hydraulic functions through biophysical feedbacks. Glob Chang Biol, 25(6), 1895–1904. doi:10.1111/gcb.14626

Rydsaa, J. H., Stordal, F., Bryn, A., & Tallaksen, L. M. (2017). Effects of shrub and tree cover increase on the near-surface atmosphere in northern Fennoscandia. Biogeosciences, 14(18), 4209–4227. doi:10.5194/bg-14-4209-2017

Scharnagl, K., Johnson, D., & Ebert-May, D. (2019). Shrub expansion and alpine plant community change: 40-year record from Niwot Ridge, Colorado. Plant Ecology & Diversity, 1–10.

Semenchuk, P. R., Elberling, B., Amtorp, C., Winkler, J., Rumpf, S., Michelsen, A., & Cooper, E. J. (2015). Deeper snow alters soil nutrient availability and leaf nutrient status in high Arctic tundra. Biogeochemistry, 124 (1-3), 81–94. doi:10.1007/s10533-015-0082-7

SFS3008. (1990). SFS 3008, Determination of total residue and total fixed residue in water, sludge and sediment. Finnish Standards Association, Helsinki, Finland, 3.

Sistla, S. A., Moore, J. C., Simpson, R. T., Gough, L., Shaver, G. R., & Schimel, J. P. (2013). Long-term warming restructures Arctic tundra without changing net soil carbon storage. Nature, 497(7451), 615-+. doi:10.1038/nature12129

Sturm, M., Holmgren, J., McFadden, J. P., Liston, G. E., Chapin III, F. S., & Racine, C. H. J. J. o. C. (2001). Snow–shrub interactions in Arctic tundra: a hypothesis with climatic implications. 14(3), 336–344.

Sturm, M., Schimel, J., Michaelson, G., Welker, J. M., Oberbauer, S. F., Liston, G. E., … Romanovsky, V. E. (2005). Winter biological processes could help convert arctic tundra to shrubland. Bioscience, 55(1), 17–26. doi:Doi 10.1641/0006-3568(2005)055[0017:Wbpchc]2.0.Co;2

Swann, A. L., Fung, I. Y., Levis, S., Bonan, G. B., & Doney, S. C. (2010). Changes in Arctic vegetation amplify high-latitude warming through the greenhouse effect. Proceedings of the National Academy of Sciences of the United States of America, 107(4), 1295–1300. doi:10.1073/pnas.0913846107

Tang, X., Li, H., Desai, A. R., Nagy, Z., Luo, J., Kolb, T. E., … Kutsch, W. J. S. r. (2014). How is water-use efficiency of terrestrial ecosystems distributed and changing on Earth?, 4, 7483.

Tarnocai, C., Canadell, J. G., Schuur, E. A. G., Kuhry, P., Mazhitova, G., & Zimov, S. (2009). Soil organic carbon pools in the northern circumpolar permafrost region.. Global Biogeochemical Cycles, 23(2). doi:Artn Gb2023 10.1029/2008gb003327

Trumbore, S. E., Chadwick, O. A., & Amundson, R. (1996). Rapid exchange between soil carbon and atmospheric carbon dioxide driven by temperature change. Science, 272(5260), 393–396. doi:DOI 10.1126/science.272.5260.393

Walker, D. A., Daniëls, F. J., Matveyeva, N. V., Šibík, J., Walker, M. D., Breen, A. L., … Hennekens, S. (2017). Circumpolar Arctic Vegetation Classification. Phytocoenologia.

Walker, M. D., Walker, D. A., Welker, J. M., Arft, A. M., Bardsley, T., Brooks, P. D., … Turner, P. L. (1999). Long-term experimental manipulation of winter snow regime and summer temperature in arctic and alpine tundra. Hydrological Processes, 13(14-15), 2315–2330. doi:Doi 10.1002/(Sici)1099-1085(199910)13:14/15<2315::Aid-Hyp888>3.0.Co;2-A

Wang, L., & Liu, H. (2006). An efficient method for identifying and filling surface depressions in digital elevation models for hydrologic analysis and modelling. International Journal of Geographical Information Science, 20(2), 193–213. doi:10.1080/13658810500433453

Williams, C. J., McNamara, J. P., & Chandler, D. G. (2009). Controls on the temporal and spatial variability of soil moisture in a mountainous landscape: the signature of snow and complex terrain. Hydrology and Earth System Sciences, 13(7), 1325–1336. doi:DOI 10.5194/hess-13-1325-2009

Wilson, S. D., & Nilsson, C. (2009). Arctic alpine vegetation change over 20 years. Global Change Biology, 15(7), 1676–1684. doi:10.1111/j.1365-2486.2009.01896.x

Winkler, M., Lamprecht, A., Steinbauer, K., Hulber, K., Theurillat, J. P., Breiner, F., … Pauli, H. (2016). The rich sides of mountain summits - a pan-European view on aspect preferences of alpine plants. Journal of Biogeography, 43(11), 2261–2273. doi:10.1111/jbi.12835

Vowles, T., & Björk, R. G. J. J. o. E. (2018). Implications of evergreen shrub expansion in the Arctic.

Vuorinen, K. E., Oksanen, L., Oksanen, T., Pyykönen, A., Olofsson, J., & Virtanen, R. J. G. c. b. (2017). Open tundra persist, but arctic features decline—Vegetation changes in the warming Fennoscandian tundra. 23(9), 3794–3807.

Ylänne, H., Olofsson, J., Oksanen, L., & Stark, S. (2018). Consequences of grazer-induced vegetation transitions on ecosystem carbon storage in the tundra. Functional Ecology, 32(4), 1091–1102.

Zevenbergen, L. W., & Thorne, C. R. (1987). Quantitative-Analysis of Land Surface-Topography. Earth Surface Processes and Landforms, 12(1), 47–56. doi:DOI 10.1002/esp.3290120107

Zwieback, S., Chang, Q., Marsh, P., & Berg, A. J. E. R. L. (2019). Shrub tundra ecohydrology: rainfall interception is a major component of the water balance. 14(5), 055005.

